# Drug repurposing approach for potential Pfmrk inhibitors as antimalarial agents: an *In-silico* analysis

**DOI:** 10.1101/2023.07.04.547694

**Authors:** Abhishek Sahu, Tanuj Handa, Debanjan Kundu

## Abstract

Malaria is a major global health issue due to the emergence of resistance to most of the available antimalarial drugs. There is an urgent need to discover new antimalarials to tackle the resistance issue. A CDK-like protein, Pfmrk from *Plasmodium falciparum*, plays a crucial role in regulating cell proliferation and shares 36.28% homology with humans CDK (hCDK7). Pfmrk complex with Pfcyc-1 and stimulates kinase activity. Also, Pfcyc-1 from P. falciparum, which has the highest sequence homology with human cyclin (Cyclin H), binds to and activates Pfmrk in a cyclin-dependent way. This is the first indication that cyclin subunits regulate human and plasmodial CDKs in a similar manner. In this study, molecular docking analysis of Pfmrk against the selected FDA-approved drugs acquired from the ZINC15 database. The top five drugs, Lurasidone, Vorapaxar, Donovex, Alvesco, and Orap, were screened based on binding energies of best-docked scores ranging between -8 kcal/mol and -12 kcal/mol. Based on Molecular dynamics simulations for 100ns, Lurasidone showed the highest binding affinity (-105.90 ± 57.72 kJ/mol), followed by Donovex (-92.877 ± 17.872 kJ/mol) and exhibited stable interactions with the amino acid residues present in the active site of Pfmrk. The outcomes of *in silico* investigation putatively suggested that Lurasidone and Donovex exhibit antimalarial potency and could be translated as potential Pfmrk inhibitors and developing new drugs based on further *in-vitro* studies.

**Graphical Abstract:** 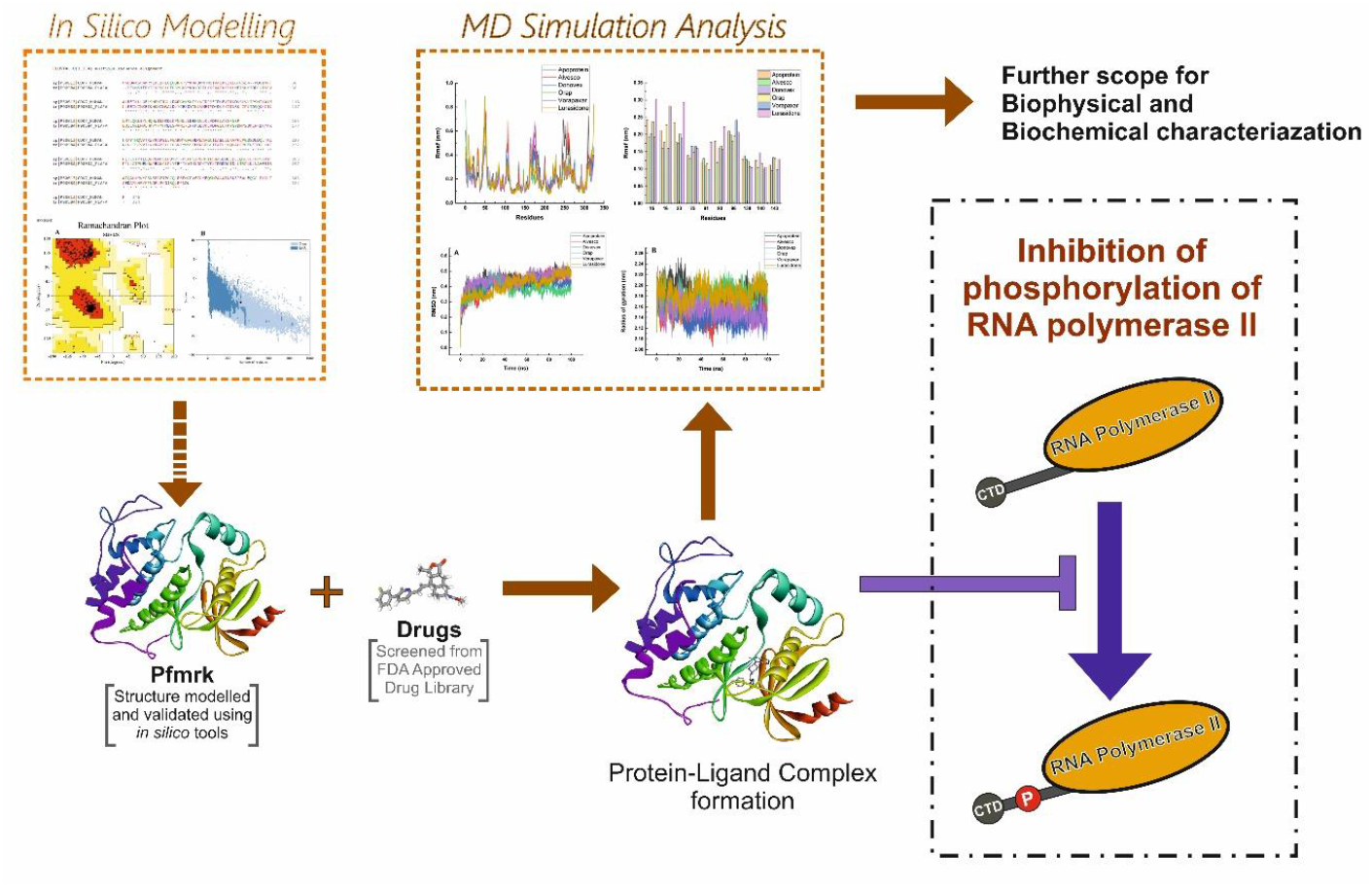

**Highlights:** We investigated the potential FDA-approved drugs for repurposing against the modelled protein Pfmrk.

Alvesco, Donovex, Lurasidone, Orap, and Vorapaxar are potential FDA-approved drugs firmly binding with Pfmrk.

Molecular docking and simulation studies show that Donovex and Lurasidone are potential inhibitors of the modelled Pfmrk protein.

Donovex and Lurasidone are potential drugs that act as kinase inhibitors by binding to the ATP -binding site of the enzyme Pfmrk.

## 1.0 Introduction

Malaria is an Italian term “mal-aria” that awful air comes from the early relation of disease with the swampy region. Malaria is generally transmitted via the bite of a female *Anopheles*-infected mosquito. Malaria is caused by a eukaryotic unicellular parasite of the genus *Plasmodium*, belonging to the Apicomplexa phylum (Tuteja *et al*., 2007). Some of the species which affect humans are *Plasmodium falciparum, P. vivax, P. ovale*, and *P. malariae*. Among these, *Plasmodium falciparum* is the fatal form of malaria. According to WHO (World Health Organization), around 300-500 million malaria cases occurred, and most cases occurred in Africa. Some patients also clustered in India, Sri Lanka, Thailand, Indonesia, Vietnam, China, and Cambodia. Out of this, more than 2 million people die because of malaria.

The parasite’s complex life cycle involves some specific protein for its continuity in vertebrate and invertebrate hosts. For their survivability in extracellular and intracellular, both these proteins are required. The parasite’s complex life cycle involves some specific protein for its continuity in vertebrate and invertebrate hosts. For their survivability in extracellular and intracellular, both these proteins are required.

Pfmrk is a member of the Apicomplexa-specific protein kinase subfamily analogous to cyclin-dependent kinases (CDK). Pfmrk, much like CDK7, behaves as an upstream kinase, thus activating multiple CDKs in a cell cycle. It is also known as CDK Activating kinase for this function. Pfmrk regulates the expression of a gene by activating the CTD of RNA Pol-II through its kinase activity, as shown in Figure 1.

**Figure 1.**
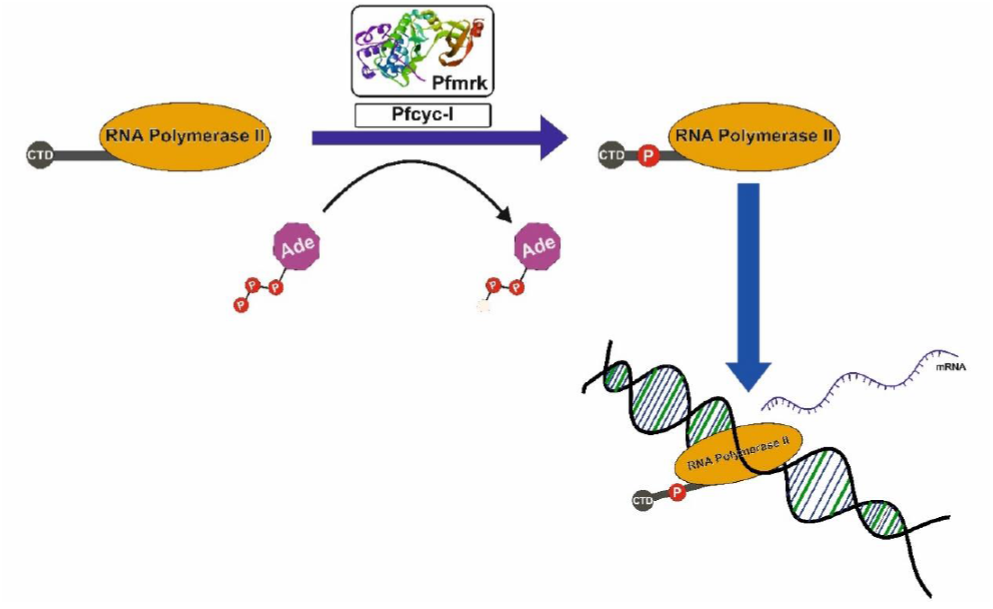
Activation of CTD of RNA Pol II by active Pfmrk-Pfcyc1 CDK complex, where CTD of RNA polymerase II acts as a substrate.

Pfmrk has dual functions in regulating gene expression and cell cycle control, which makes Pfmrk a potential drug target. Pfcyc-1 from *P. falciparum* shares maximum sequence homology with human cyclin (Cyclin H), associated with and activating Pfmrk in a cyclin-dependent manner. Several studies revealed that the C-terminal domain of RNA Pol II is an endogenous substrate of Pfmrk (Keenan *et al*., 2005). Thus Pfmrk is chosen to be a potential target for antimalarial drug screening.

## 2.0 Methodology

### 2.1 Sequence homology of Pfmrk with Hcdk7

The sequence homology of Pfmrk with Hcdk7 was performed using multiple sequence alignment through Clustal Omega (1.2.4) tool (McWilliam *et al*., 2013). The FASTA sequence of Pfmrk (ID: P90584) and hCDK7 (ID: P50613) was obtained from the UniProt database (UniProt Consortium, 2019). Further, the clustal omega tool was used to estimate the sequence identity between Pfmrk and hCDK7, as shown in Figure S1.

### 2.2 Protein Preparation

In the absence of the experimental structure of target protein Pfmrk, the ‘FASTA sequence of target enzymes Pfmrk (ID: P90584) was retrieved from the UniProt database. The Galaxy TBM server (Ko *et al*., 2012) was used to build the protein model, as shown in Figure 2. The validation of a model of the model was done using the SAVES v6.0 server PROCHECK (Laskowski *et al*., 2001) for Ramachandran plot analysis. The model with more significant % residues in the most favored region was selected (Fig S2.a). Further, the quality of the model was validated using the PROSa webserver (Fig S2.b). The energy minimization of the modeled protein was done using SWISS PDB viewer software (Geux *et al*., 1997).

**Figure 2.**
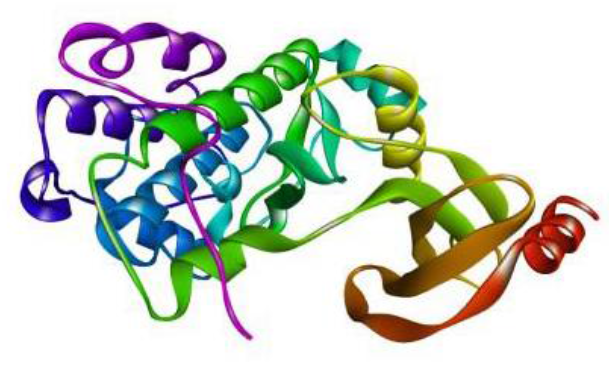
Structure of Pfmrk obtained through Galaxy TBM Web server.

### 2.3 Active site identification & selection for molecular docking

The active site was chosen based on the literature that is currently available on the catalytically active protein Pfmrk sites. The active site of Pfmrk contains residues such as L16, M75, M91, I93, & F143, which form a hydrophobic pocket for binding the ATP. However, the ATP binding site and these five residues in hCDK7 are replaced by different amino acids, such as D16, I75, F91, F93, and L143(Waters *et al*., 2000). The grid box was prepared along these five residues lying in the active site of Pfmrk. The presence of amino acids in the docking site of protein with detailed coordinates is shown in Table 1.

**Table 1.** Shows information about the docking proteins, the docking sites, and the relevant coordinates and references.

| Protein | Docking site | Coordinates (X, Y, Z) | Reference |
| --- | --- | --- | --- |
| **Pfmrk** | Phe15, Lys23, Met91, Ile93, Tyr96, Ser138, Ala140, Phe143, | X_266.09, Y_319.852, Z_310.061 | Keenan et al. 2005 |
|  | **ATP binding site** |  |  |
|  | Leu16, Met75, Met91, Ile93, Phe143 | X_266.09, Y_319.852, Z_310.061 | Waters et al, 2000 |

### 2.4 Ligand Preparation

The 1576 FDA-for-sale compounds from the ZINC 15 database in MOL2 format (https://zinc.docking.org/substances/subsets/fda+for-sale/). According to Lipinski’s rule, the duplicates, empty structures (salts), isotopes, and inorganics were screened and removed using the FAF drugs4 server (Lagorce *et al*., 2008). The final database obtained contained 1467 compounds. All the compounds from ZINC15 are in ready-to-dock conformations of the compounds. All the compounds in MOL2 format were imported to PyRx software for energy minimization and were converted to PDBQT format for docking (Dallakyan *et al*., 2015).

### 2.5 Molecular docking and compound screening

The MGLTools v.1.5.6 Autodock was used for molecular docking using Raccoon Virtual Screening (Morris *et al*., 2009). The grid was centered around the active site of Pfmrk containing amino acid residues, and the grid box was constructed with coordinates dimensions: center_ x = 266.09, center_y = 319.852, center_z = 310.061, size_x = 60, size_y = 54, size_z = 54. Tools v.1.5.6 Autodock was used for molecular docking using Raccoon Virtual Screening (Morris *et al*., 2009). The principal docking protocol employs AUTODOCK4 and the common Lamarckian Genetic Algorithm (LGA) (Morris et al., 1996; Kundu et al., 2020). The best compound was selected based on the lowest binding free energy (kcal/mol) and the combination with the highest number of clusters combination present.

### 2.6 Molecular dynamic simulation research

Studies on molecular dynamic simulations support docking results for apoprotein and protein-ligand complexes. Using GROMACS v 2018.8, the top five drugs were simulated as a positive control. Using the GROMOS force field, PRODRG server version 2.5 (Borkotoky et al., 2021; Van et al., 1996) produces boundary and topographic files for ligands. The GROMOS 54a7 force field was deployed to prepare the apoprotein and ligand complexes, and SPC/E water molecules were used to add solvation to our system in a cubic frame with a dimension of 1.2 nm (Abraham et al., 2015).

The system was neutralized by adding the appropriate number of Na+ and Cl-ions during the ionization step. For all the systems, the energy minimization step was carried out with an acceptance of 1000 KJ/mol, and a maximum number of steps was set at 50,000. The PME method was deployed to establish the cutoff value of 1.2 nm for both long-range and short-range interactions. Further, the NVT and NPT equilibration procedure was performed for 1ns with a fixed particle number, volume, and temperature during the post-energy minimization. NVT equilibration was done using the Berendsen thermostat (Borkotoky *et al*., 2021; Bussi *et al*., 2007) with a pace of 0.002 fs at a temperature of 300 K. To identify long-range interactions, the Particle Mesh Ewald (PME) method (Kawata & Nagashima, 2001) was used, with a cutoff value of 1.2 nm and 0.16 nm set for Fourier spacing. Also, for NPT equilibration, several particles, pressure, and temperature were fixed. To achieve NPT equilibration, Berendsen isotropic pressure, a time constant of 2 fs, a pressure bar of 1, and isothermal compaction of 4.5 e-5 bar are all required. Following the conclusion of both equilibration steps, simulations of each system were run for 100 ns with a dt of 2 fs and a leap-frog integrator (Borkotoky et al., 2021). Finally, the simulation results of all the systems were analyzed using the standard commands through the GROMACS platform, and the LINCS algorithm was used to restrain all bond lengths (Hess *et al*., 1997; Kundu & Dubey, 2021).

### 2.7 MM/PBSA analysis

The MM/PBSA method was performed using the gmx mmpbsa tool and the pbsa.mdp script, which evaluates the free binding energy of the top five drugs to the apoprotein and their binding mode. To save computation time, the free energy evaluation was carried out for the final 40ns of the run. The value of dielectric constants for solute and solvent were 2 and 80, respectively (Genheden & Ryde, 2015; Musyoka et al., 2016; Kundu & Dubey, 2021).

### 2.8 Visual analysis

The Discovery Studio Visualizer software was used to generate a two-dimensional diagram of the protein-ligand complex possible interactions for further study (Dassault Systemes BIOVIA 2020). Potential hydrophobic interactions were analyzed using Protein-ligand Interaction Profiler (Salentin et al., 2016; Kundu & Dubey, 2021).

## 3.0 Results

### 3.1 Docking results

As previously mentioned, Pfmrk active site includes non-conserved amino acids such as Phe15, Leu16, Lys23, Met75, Met91, ILe93, Tyr96, Ser138, Ala140, and Phe143. The maximum number of amino acids was enclosed within the grid box. The top five drugs named, Lurasidone, Discovery studio visualizer Vorapaxar, Donovex, Alvesco, and Orap, were selected having lowest binding energy and show the maximum possible interaction with amino acids from the enzyme’s active site (Figure 3).

**Figure 3.**
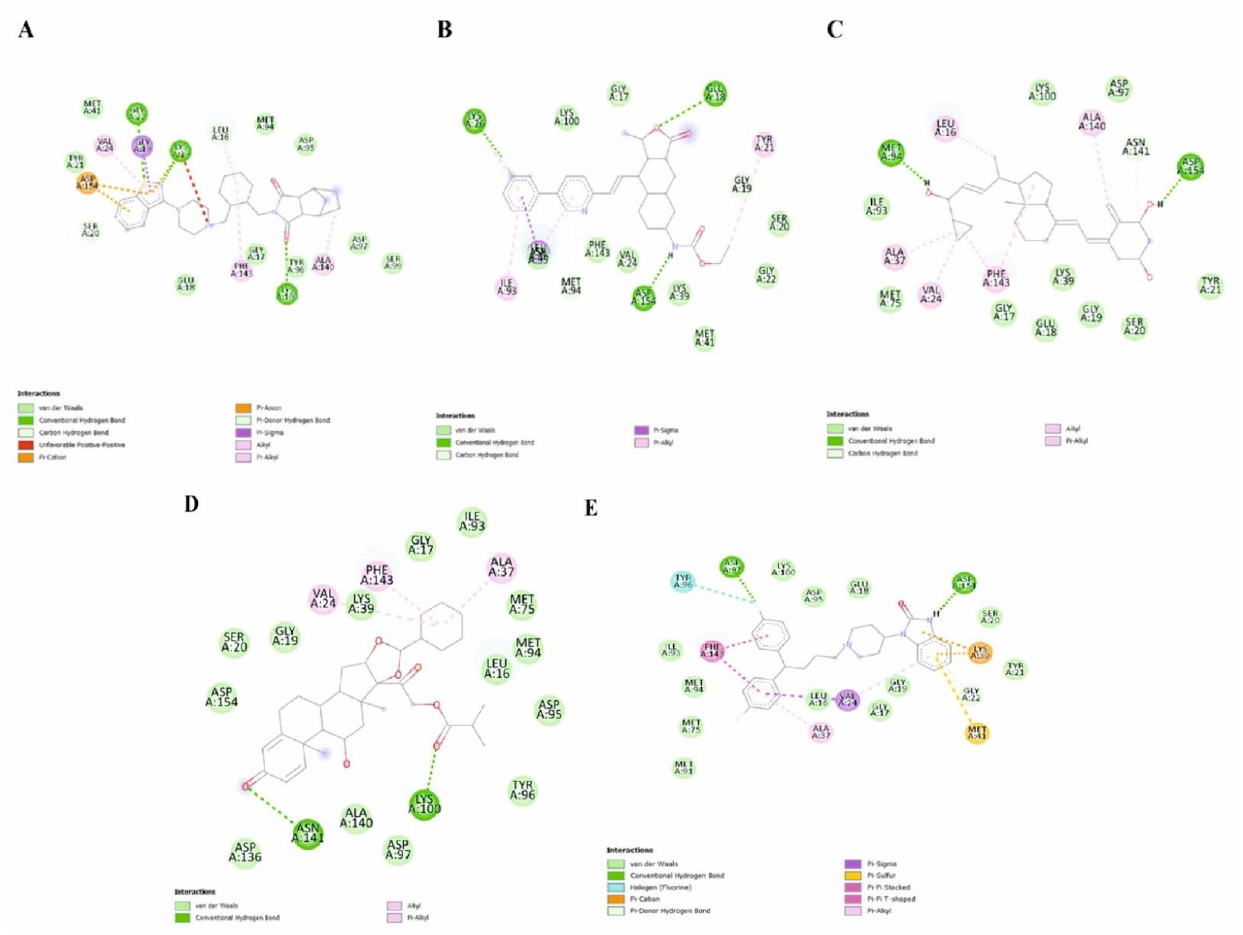
2D Interaction of Pfmrk with (A) Lurasidone, (B) Vorapaxar, (C) Donovex, (D) Alvesco, and (E) Orap as visualized in Biovia Discovery studio visualizer

Among these top five drugs, Lurasidone was found to have the lowest binding energy (-12.03 kcal/mol) with Ki of 1.52nM. Lurasidone forms a non-covalent interaction (alkyl and pi-alkyl) with Ala140 & Phe143, which are critical residues of the PAS of the enzyme. Vorapaxar had binding energy of (-10.06 kcal/mol) with the Ki of 39.14nM and exhibited non-covalent interaction (pi-alkyl) with the Ile93, an essential residue of the enzyme. Besides these two drugs, Donovex, Alvesco, and Orap exhibit higher binding energy with higher Ki value and show non-covalent interactions (alkyl and pi-alkyl) with Phe143 and Ala140 key residues of PAS of the enzyme. Also, a significant non-covalent interaction was observed between apoprotein and drug in the ATP-binding region and the area around our ligand-binding site. The details of the top five drug interactions, binding free energy, and inhibitor constant (Ki) as determined by molecular docking are shown in Table 2.

**Table 2.** Shows various kinds of FDA drugs interactions, binding free energy and inhibitor

| **FDA Drug** | **Free Binding energy (kcal/mol)** | **Ki nanomole** | **Hydrogen bond Interactions** | **Non-covalent bond Interactions** |
| --- | --- | --- | --- | --- |
| **Lurasidone** | -12.03 | 1.52nM | Lys39, Lys100, Gly22 | Leu16, Val24, Phe143 pi-alkyl, Ala140, Asp154(2), lys39pi-anion, Gly19 pi-sigma, |
| **Vorapaxar** | -10.06 | 39.14nM | Lys26, Glu18, Asp15, | Ile93 Pi-alkyl, Tyr21, Gly19C-H bond, Met94, Leu16pi-sigma |
| **Donovex** | -8.69 | 429.05nM | Met94(2), Asp154 | Leu16, Ala140, Phe143(2), Val24, Ala37 |
| **Alvesco** | -8.68 | 435.75nM | Asn141, Lys100 | Val24(2), Ala37, Ala140, Phe143 |
| **Orap** | -8.46 | 633.3nM | Asp154, Asp97 | Lys39(2), Phe143(2) Pi-Pi stacked, Ala37 pi alkyl, Met41 Pi cation, Val24, Lys39 |

### 3.2 Molecular simulation

The Pfmrk apoprotein and its complexed ligands were simulated for 100 ns. Five different types of analysis were performed on simulated complexes to support docking results. The RMSD curve was obtained for both the apoprotein and holoprotein clusters using the gmx_rms tool, which indicates the systems’ overall stability. The gmx gyrate tool was used throughout the simulation to calculate the system’s gyration radius. The tool gmx rmsf was used to calculate the rmsf for the total protein and complex. The mean number of hydrogen bonds between the apoprotein and complex was calculated using the gmx hbond tool to assess their stability further. An atomic distance cutoff of 0.35 nm was set between the donors and acceptors.

### 3.3 RMSD, Radius of Gyration and RMSF

The RMSD curve of apoprotein and the other five complexes were compared and reported that the vorapaxar curve is similar to that of the apoprotein. All five drug complexes show minimal fluctuations, and the curve of all the drugs converges between 0.3 to 0.5nm. Alvesco and Donovex shows excellent stability over the entire simulation run of 100ns. The Orap complex shows sudden fluctuations up to 40ns, and RMSD was stable afterward Figure 4(A). All five complexes have a greater degree of compactness than the apoprotein overall in terms of the radius of the gyration curve (Rg). Donovex, Vorapaxar, and Alvesco were found to have the highest degree of compactness, with the highest deviation of 2.12nm. Orap and Lurasidone showed a curve like an apoprotein, which converges around 2.22nm Figure 4(B).

**Figure 4.**
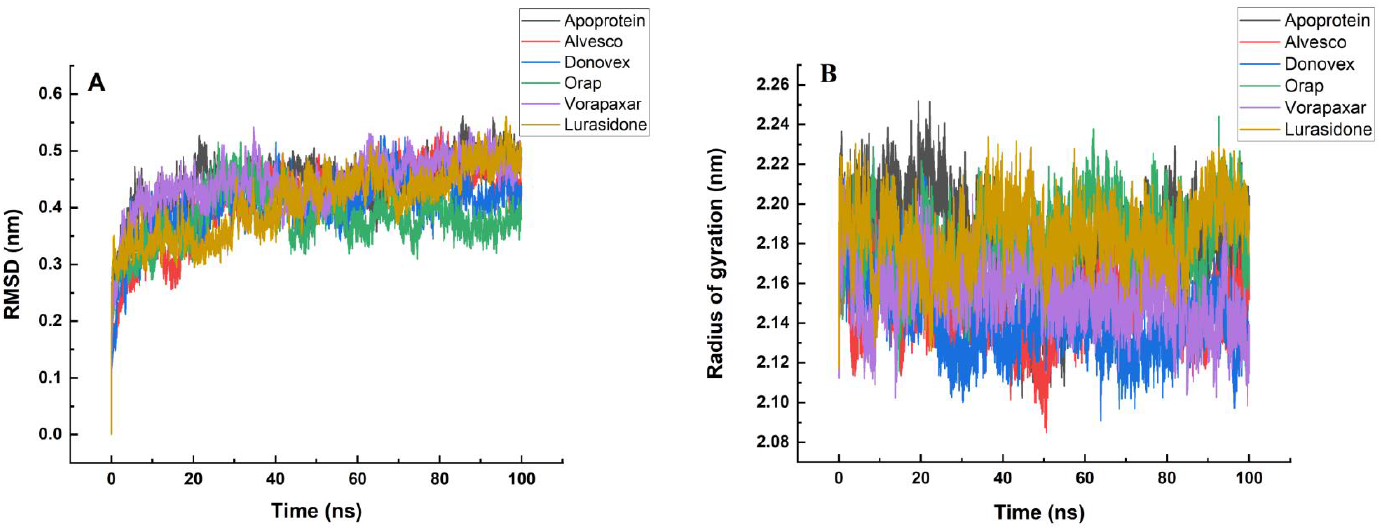
Analysis of the RMSD and Rg results from an MD simulation. A. Comparison of the RMSD trajectories for 100 ns of all protein-ligand complexes and apoproteins B. A comparison of the Rg values for each protein-ligand complex and the apoprotein over a 100 ns period.

As per the docking result, Alvesco and Donovex found a Non-covalent interaction with the Ala140, Phe143, and Leu 16 residues present in the active site of the apoprotein, consequently reducing the RMSF curve between residues 15-143. At the same time, Lurasidone shows a more excellent RMSF curve trajectory between the entire amino acids stretch of 15-143 (Figure 5).

**Figure 5.**
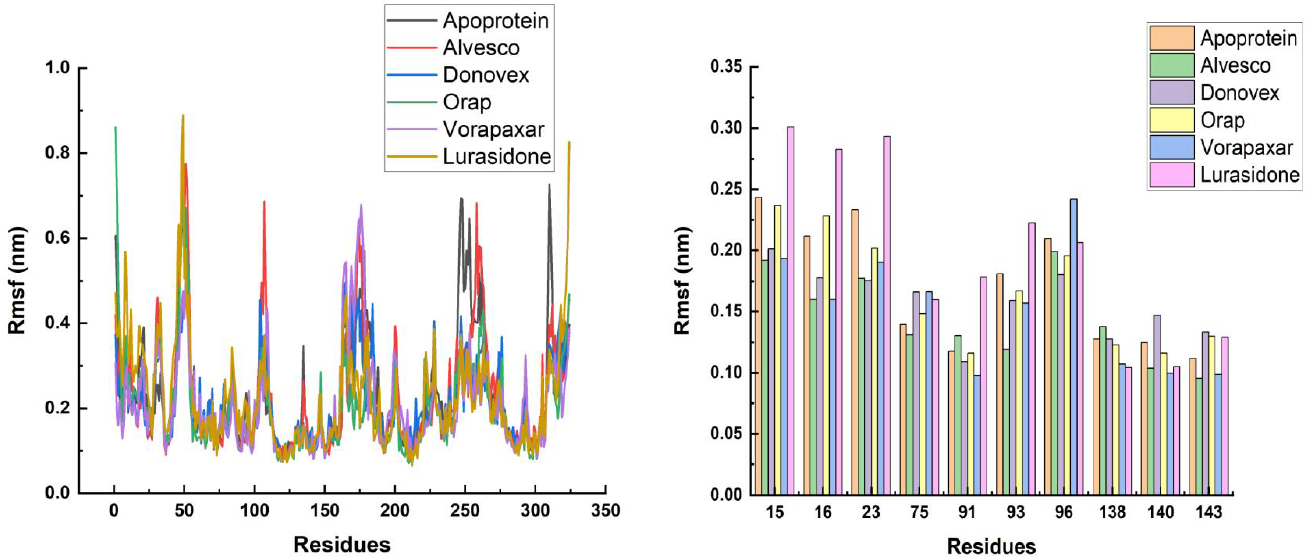
Analysis of the RMSF results from an MD simulation. A comparison of all six drug complexes and the apoprotein RMSF values for a 100 ns course B. Comparing the RMSF values for the active site of each of the six drug complexes and the apoprotein for a 100 ns course.

### 3.4 SASA and Hydrogen Bonding

The Solvent Accessible Surface Area (SASA) result of all protein-ligand complexes was lower in value than apoprotein, indicating a strong binding affinity between ligands and apoprotein and increasing the apoprotein’s compactness. The SASA result of all protein-ligand complexes is represented in **Table S1**. Simultaneously, the apoprotein was found to be more folded in the presence of ligands. The apoprotein complexed with Donovex was found to have the lowest SASA value, thus increasing the compactness of the apoprotein. The H-bond analysis of all protein-ligand complexes is represented in **Table S2**. H-bond analysis using MD simulation shows that Alvesco forms three hydrogen bonds with Lys100, Ser138 & Ala140 and forms one average hydrogen bond with the amino acid residues present in the active site of the apoprotein.

In contrast, the docking result of Alvesco also constitutes one h-bond, Lys100) in static conditions shown in our docking result. Donovex forms more hydrogen bonds with amino acids (Met 94, Asp154, Ile 93, Ser138, Ala140 & Phe143) and includes one average h-bond with residues present in the active site of our apoprotein. In contrast, the docking result of Donovex also forms a two h-bond with (Met 94 & Asp 154). Lurasidone forms hydrogen bonds with amino acids (Gly22, Lys39, Lys100). It includes one average h-bond with residues present in the active site of our apoprotein, whereas docking results in Lurasidone also forms a three h-bond with (Gly22, Lys39, Lys100). Our results indicate that the hydrogen bonds depicted during our docking are often unstable over a long simulation period. Stable hydrogen bonds are necessary for protein-ligand solid interactions.

Orap forms two hydrogen bonds with amino acids (Lys23 & Asp154) and forms two average h-bonds with residues present in the active site of our apoprotein. In contrast, the docking result of Orap also forms a one h-bond with Asp154). In comparison, Vorapaxar forms one hydrogen bond with amino acids (Glu18, Lys26, Tyr96, Lys100) residues in our binding site in contrast to our docking result where it formed two hydrogen bonds with Lys 26 & Glu18.

### 3.5 Secondary structure analysis

The comparative DSSP secondary structure analysis of both apoprotein and ligands was calculated using MD simulation and is represented in **Table S3**. Secondary structure analysis of apoprotein shows a similar content of β-sheet, β-bridge, bend, turn, α-helix, 5-helix & 3-helix with ligands. Also, Alvesco and Donovex offer a higher range of (coil, β-sheet, β-bridge, and bend), suggesting that ligands have maximum non-covalent interaction with the apoprotein. Orap shows a greater content of turn, meaning it contributes more significantly toward protein compactness than others.

### 3.6 MM PBSA free energy analysis

The overall free binding energy for Donovex was found to be -92.87+/-17.87 kJ/mol and Lurasidone was -105.9+/-57.72 kJ/mol, which was higher as compared to other ligands, as shown in Table3. Donovex and Lurasidone are shown to have the highest contributor to Van der Waals forces, the significance of hydrophobic interactions in our protein-ligand interactions, and the influence the ligands have on a more folded state of the apoprotein. Donovex and Lurasidone bind to the protein’s active site and other binding compartments with relatively high binding affinity than other ligands, according to the noticeably low binding energy.

**Table 3.** Detailed comparative analyses of all the energetic components of the protein-drug complexes formed post-simulation.

| LIGANDS | Van der Waal energy KJmol<sup>-1</sup> | Electrostatic energy (KJmol<sup>-1</sup>) | Energy of solvation (KJmol<sup>-1</sup>) | SASA energy (KJmol<sup>-1</sup>) | Binding energy (KJmol<sup>-1</sup>) |
| --- | --- | --- | --- | --- | --- |
| Alvesco | -121.789 ± 69.283 | -1.518 ± 7.973 | 65.713 ± 46.072 | -13.350 ± 7.725 | -70.944 ± 54.776 |
| Donovex | -126.353 ± 20.061 | -2.796 ± 2.679 | 51.327 ± 15.995 | -15.054 ± 2.503 | -92.877 ± 17.872 |
| Lurasidone | -127.589 ± 64.217 | -7.726 ± 7.060 | 40.816 ± 23.777 | -11.408 ± 5.727 | -105.907 ± 57.728 |
| Orap | -103.655±100.467 | -3.910 ± 6.977 | 42.897 ± 56.765 | -9.701 ± 9.853 | -74.369 ± 75.259 |
| Vorapaxar | -5.003 ± 28.557 | -0.642 ± 4.782 | 16.218 ± 40.884 | -0.269 ± 4.157 | 10.304 ± 41.271 |

## 4.0 Discussion & Conclusion

This computational work aims to identify the potential FDA-approved drugs against Pfmrk as antimalarial drugs. Molecular docking with FDA-approved drugs and molecular dynamics simulation was performed to confirm molecular docking findings. As discussed above, the active site of Pfmrk contains residues such as Leu16, Met75, Met91, Ile93, & Phe143, forming a hydrophobic pocket for binding the ATP, and these five residues are substituted by different amino acids Asp16, Ile75, Phe91, Phe93, Leu143 in hCDK7. Based on the molecular docking results, it was inferred that hydrogen bonds, hydrophobic and other non-covalent interactions are critical players for strong binding between apoprotein and ligand. Given that Pfmrk has dual functions in cellular replication and the overall life cycle of *P. falciparum*, inhibiting the binding of ATP by other compounds could be an essential strategy.

The current study aimed to screen and validate some drugs that could inhibit Pfmrk kinase activity by strongly binding to the protein’s ATP binding site.

The top five FDA drugs obtained through docking results were selected based on the higher binding affinity and lower energy. The top five FDA-approved drugs, such as Lurasidone, Vorapaxar, Donovex, Alvesco, and Orap, all these drugs’ original use and year of acceptance were reported in **Table S4**.

Among these five drugs, it was found that Donovex and Lurasidone have the lowest binding free energy of -8.69 kcal/mol and -12.03 kcal/mol through molecular docking and have non-covalent interactions with Leu16, Ala140 & Phe 143. Furthermore, some critical key residues, such as Ile93 and Phe143, are the key residues in Pfmrk’s ATP binding site with which ligands interact. Phe143 shows varied non-covalent interactions like Pi-alkyl, whereas Ile93 forms Van-Der Waal’s interactions. The significance of molecular dynamics and simulation is that it supports molecular docking analysis. The MD simulation analysis confirms that Donovex and Lurasidone bind with the ATP binding site with greater binding affinity, as presented in our RMSD, RMSF, and the radius of gyration analyses. The other research, such as SASA, H-bond, and MM-PBSA analysis, also shows a better impact of Donovex and Lurasidone on the target protein. The interaction of Lurasidone and Donovex with Phe143 & Ile93, which are essential residues in the ATP binding region, coupled with low binding energy and other favorable results, indicate that these ligands in the ATP binding region might prove to inhibit the binding of ATP, leading to inhibition of kinase activity of Pfmrk. Further, the inactivation of CTD of RNA Pol II stops the transcription machinery from working.

As per the research done till now, it was found that *P. falciparum* parasites are strongly resistant to antimalarial drugs, which led to identifying new targets. Protein kinase such as Plasmodial CDK (Pfmrk) satisfies all the conditions above for a promising drug target. Sequence homology between Pfmrk and hCDK7 in the current study showed the sequence identity to be 36.28%, suggesting a considerable difference between the two proteins. The significant differences between Pfmrk and hCDK7 suggest drug for inhibiting this enzyme will not cause any harm to the host itself.

## Acknowledgment

We acknowledge research fellowships by IIT (BHU) Varanasi. The infrastructure support and the resources provided by DST funded I-DAPT Hub Foundation, IIT BHU [DST/NMICPS/TIH11/IIT(BHU)2020/ 02] and PARAM Shivay Facility under the National Supercomputing Mission, Government of India at the Indian Institute of Technology, Varanasi are gratefully acknowledged. AS, TH, and DK acknowledges the research fellowship provided by IIT (BHU).

