## Supplementary materials for "Drug repurposing approach for potential Pfmrk inhibitors as antimalarial agents: an *In-silico* analysis"

```
CLUSTAL O(1.2.4) multiple sequence alignment


sp|P50613|CDK7_HUMAN        MALDVKSRAKRYEKLDFLGEGQFATVYKARDKNTNQIVAIKKIKLGHRSEA--KDGINRT    58
tr|P90584|P90584_PLAFA      --MENNSTERYIFKPNFLGEGSYGKVYKAYDTILKKEVAIKKMKLNEISNYIDDCGINFV    58
                              :: :*  :   * :*****.:..**** *.  :: *****:**.. *:   . *** .

sp|P50613|CDK7_HUMAN        ALREIKLLQELSHPNIIGLLDAFGHKSNISLVFDFMETDLEVIIKDNSLVLTPSHIKAYM    118
tr|P90584|P90584_PLAFA      LLREIKIMKEIKHKNIMSALDLYCEKDYINLVMEIMDYDLSKIINRK-IFLTDSQKKCIL    117
                             *****:::*:.* **:. ** : .*. *.**:::*: **. **: : :.** *: *. :

sp|P50613|CDK7_HUMAN        LMTLQGLEYLHQHWILHRDLKPNNLLLDENGVLKLADFGLAKSFGSP-------------    165
tr|P90584|P90584_PLAFA      LQILNGLNVLHKYYFMHRDLSPANIFINKKGEVKLADFGLCTKYGYDMYSDKLFRDKYKK    177
                            *  *:**: **:::::****.* *::::::* :*******...:*

sp|P50613|CDK7_HUMAN        NRAYTHQVVTRWYRAPELLFGARMYGVGVDMWAVGCILAELLLRVPFLPGDSDLDQLTRI    225
tr|P90584|P90584_PLAFA      NLNLTSKVVTLWYRAPELLLGSNKYNSSIDMWSFGCIFAELLLQKALFPGENEIDQLGKI    237
                            *   * :*** ********:*:. *. .:***:.***:*****:  ::**:.::*** :*

sp|P50613|CDK7_HUMAN        FETLGTPTEEQWPDMCSLPDYVTFKSFPGIPLHHIFSAAGDDLLDLIQGLFLFNPCARIT    285
tr|P90584|P90584_PLAFA      FFLLGTPNENNWPEALCLPLYTEFTKATKKDFKTYFKIDDDDCIDLLTSFLKLNAHERIS    297
                            *  ****.*::**:  .** *. *..     ::  *.  .** :**: .:: :*   **:

sp|P50613|CDK7_HUMAN        ATQALKMKYFSNRPGPTPGCQLPRPNCPVETLKEQSNPALAIKRKRTEALEQGGLPKKLI    345
tr|P90584|P90584_PLAFA      AEDAMKHRYFFNDPLPCDISQLPFNDL---------------------------------    324
                            * :*:* :** * * *   .***  :

sp|P50613|CDK7_HUMAN        F    346
tr|P90584|P90584_PLAFA      -    324
```

Figure S1. Showing Multiple sequence alignment of Pfmrk & hCDK7 using CLUSTAL O ((1.2.4) and **Sequence identity- 36.28%.**

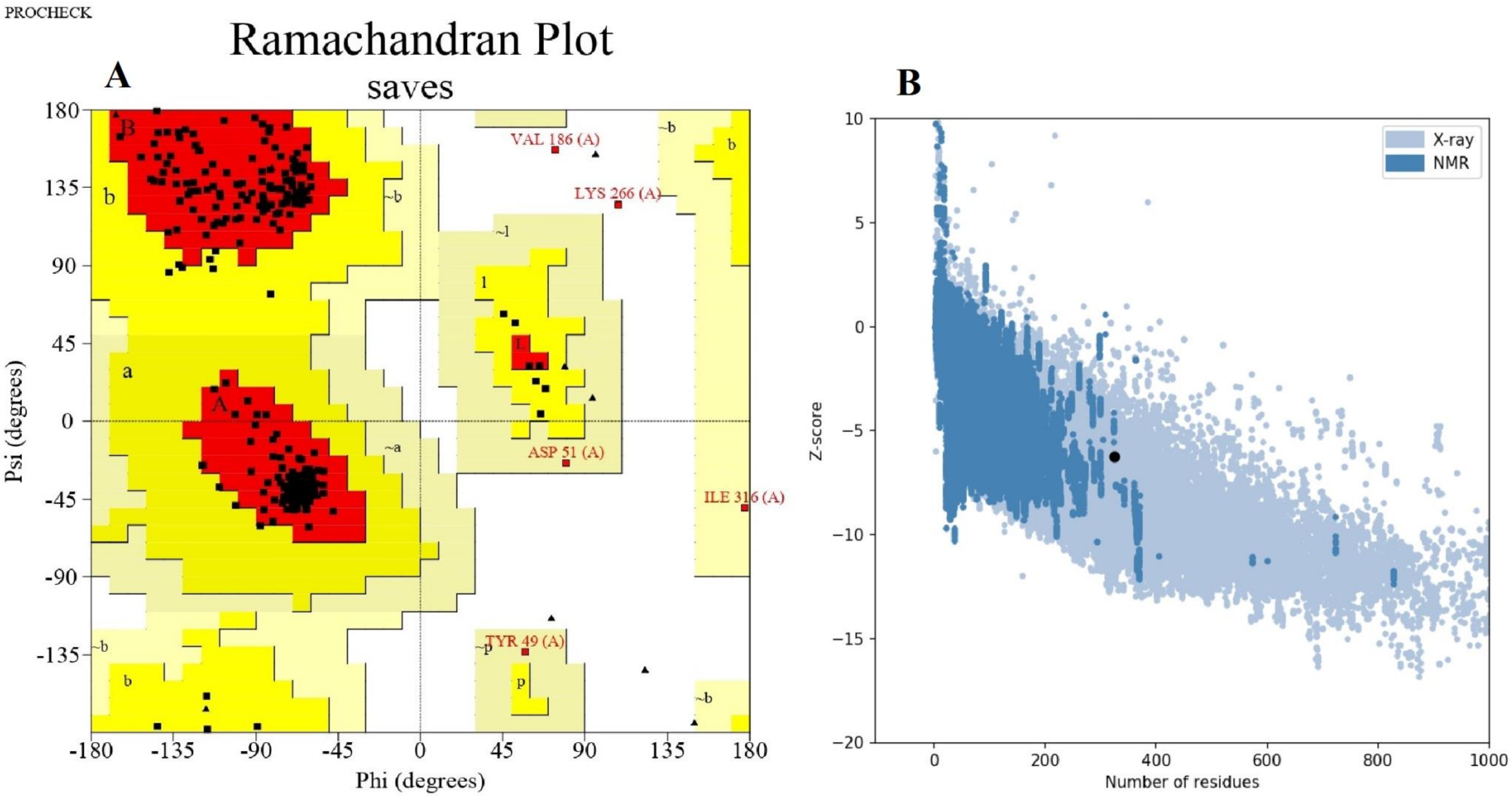


Figure S2. A Ramachandran plot for the modeled structure of Pfmrk was checked through Procheck. Residues in most favored regions are 92.3%, and residues in disallowed regions are 0.7%. b Q mean score for the modelled structure of Pfmrk through ProSA web is Z-score -6.25.

Table S1. Solvent Accessible Surface Area of model protein in apostate and in complex with various ligands

| **MODEL** | **SASA($nm^2$)** |
|---|---|
| Apoprotein | 184.0569 |
| Alvesco | 181.9433 |
| Donovex | 178.1943 |
| Lurasidone | 181.3419 |
| Orap | 183.1197 |
| Vorapaxar | 180.9219 |

Table S2. Hydrogen bond analysis between apoprotein and various ligands used in the study

| **LIGANDS** | **H- Bonding** | **Average H- bonds** |
|---|---|---|
| **Alvesco** | Lys100, Ser138, Ala140 | ~1 |
| **Donovex** | Met94, Asp154, Ile 93, Ser138, Ala140, Phe143 | ~1 |
| **Lurasidone** | Gly22, Lys39, Lys100 | ~1 |
| **Orap** | Lys23, Asp154 | ~2 |
| **Vorapaxar** | Glu18, Lys26, Tyr96, Lys100 | ~1 |

Table S3. Secondary Structure Analysis of apoprotein and apoprotein in various complexes with different ligands

| **LIGANDS** | **Coil (%)** | **B-Sheet (%)** | **B-Bridge (%)** | **Bend (%)** | **Turn (%)** | **A- Helix (%)** | **5- Helix (%)** | **3- Helix (%)** |
|---|---|---|---|---|---|---|---|---|
| Apoprotein | 28 | 13 | 1 | 15 | 10 | 31 | 0 | 3 |
| Alvesco | 29 | 12 | 1 | 15 | 12 | 27 | 2 | 1 |
| Donovex | 27 | 12 | 1 | 16 | 12 | 27 | 2 | 2 |
| Lurasidone | 28 | 13 | 1 | 14 | 11 | 32 | 0 | 3 |
| Orap | 27 | 13 | 1 | 13 | 13 | 31 | 0 | 2 |
| Vorapaxar | 28 | 12 | 2 | 15 | 12 | 28 | 3 | 1 |

Table S4. FDA Approved drug's original use and year of Approval

| **Sr. No** | **Drug Approved** | **Original Use** | **Year of Approval** |
|---|---|---|---|
| 1 | Lurasidone | Schizophrenia and Bipolar I disorder | 2010 |
| 2 | Vorapaxar | Reduce Myocardial infarction | 2014 |
| 3 | Donovex | Psoriasis | 1999 |
| 4 | Alvesco | Symptomatic relief of nasal symptoms | 2008 |
| 5 | Orap | Tourette's Disorder | 1985 |
